# CRISPRi-based functional genomic screening identifies genes essential for CH_4_-dependent growth in the methanotroph *Methylococcus capsulatus* Bath

**DOI:** 10.1101/2025.05.28.656465

**Authors:** Jessica M. Henard, Spencer A. Lee, Yao-Chuan Yu, Danyang Shao, Rajeev K. Azad, Calvin A. Henard

**Affiliations:** Department of Biological Sciences and BioDiscovery Institute, University of North Texas, Denton, TX, USA

**Keywords:** methanotroph, CRISPRi, methane monooxygenase, RubisCO, functional genomics

## Abstract

Aerobic methanotrophic bacteria are the primary organisms that consume atmospheric methane (CH_4_) and have potential to mitigate the climate-active gas. However, a limited understanding of the genetic determinants of methanotrophy hinders the development of biotechnologies leveraging these unique microbes. Here, we developed and optimized a CRISPR interference (CRISPRi) system to enable functional genomic screening in methanotrophic bacteria. We built a genome-wide single guide RNA (sgRNA) library in the industrial methanotroph, *Methylococcus capsulatus*, consisting of ∼45,000 unique sgRNAs mediating inducible, CRISPRi-dependent transcriptional repression. A selective screen during growth on CH_4_ identified 233 genes whose transcription repression resulted in a fitness defect and repression of 13 genes associated with a fitness advantage. Enrichment analysis of the 233 putative essential genes linked many of the encoded proteins with critical cellular processes like ribosome biosynthesis, translation, transcription, and other central biosynthetic metabolism, highlighting the utility of CRISPRi for functional genetic screening in methanotrophs. The CRISPRi screen uncovered that 50 ppb Co^2+^ in methanotroph Nitrate Mineral Salts medium is growth inhibitory and decreasing Co^2+^ to < 10 ppb improved biomass productivity from CH_4_. Collectively, our results show that the CRISPRi system and sgRNA library developed here can be used for facile gene-function analyses and genomic screening to identify novel genetic determinants of methanotrophy and isolation of improved methanotroph biocatalysts.

## Introduction

Aerobic methanotrophic bacteria, methanotrophs, play an essential role in balancing atmospheric carbon by oxidizing methane (CH_4_) to multi-carbon molecules (methanotroph biomass) and carbon dioxide (CO_2_). These ubiquitous microbes limit the escape of CH_4_ to the atmosphere via biofiltration in terrestrial and aquatic systems^1^. Methanotrophs can also be leveraged for the bioconversion of abundant CH_4_-rich gas streams from oil and natural gas extraction sites, coal mines, agricultural waste, landfills, and wastewater treatment facilities, and have the potential to capture rising CH_4_ greenhouse gas and mitigate climate change^2^. However, significant advances in methanotroph biology and engineering for effective CH_4_ capture and conversion are needed to realize cost-effective solutions using methanotrophs.

Methanotrophs belong to a broad group of methylotrophic microbes that use organic single carbon (C_1_) molecules as sources of carbon and/or energy, which, in addition to CH_4_, includes methanol (CH_3_OH), formate (CH_2_O), and/or methylamine. Unique to methanotrophs is the CH_4_ monooxygenase (MMO) enzyme that oxidizes the strong C-H bond of CH_4_ to generate CH_3_OH^3^. Methanotroph C_1_ metabolic enzymes and pathways for CH_3_OH assimilation to biomass and its dissimilation to CO_2_ generally overlap with those in methylotrophs^4^. These include the ribulose monophosphate and serine cycles used by Gammaproteobacterial methanotrophs and Alphaproteobacterial methanotrophs for C_1_ assimilation, respectively^5–7^. In addition, the methanotroph dissimilatory metabolic processes that couple C_1_ oxidation to the electron transport chain for ATP production are coincident with those in other methylotrophs^89^. These central CH_4_ metabolic pathways have been biochemically validated using radiolabeled isotopes in a subset of methanotrophs^5,7,10^ and the availability of several complete methanotroph genomes and comparative genomics have provided critical insight into methanotroph CH_4_ metabolism^11–18^. However, many genes that may be required for methanotrophy, especially those outside of the core C_1_ metabolic gene clusters, are difficult to identify due to their strict CH_4_-dependent cultivation.

Early research using chemical and insertional transposon mutagenesis of the CH_3_OH-utilizing methylotrophic bacterium *Methylobacterium extorquens* AM1 identified many genes required for methylotrophic growth, including those encoding the enzymes needed for methanol oxidation, formaldehyde oxidation, formate oxidation, and assimilation via the serine cycle^6,19–26^. Additionally, transposon mutagenesis led to the identification of genes required for transport and utilization of rare earth elements^27^ and transcriptional regulators of the C_1_ metabolic pathways^20,28^. Notably, the elucidation of genes required for methylotrophy was enabled by leveraging their facultative capacity to utilize alternative multi-carbon sources (i.e. succinate) supplied during library generation^28^. In contrast, the vast majority of methanotrophs are obligate CH_4_-utilizers; thus, mutations in essential genes required for growth on CH_4_ would be lost from the mutagenized population. There are only two reports of successful chemical mutagenesis^29^ and one report of transposon insertional mutagenesis^30^ in methanotrophs, underscoring barriers applying forward genetic techniques to CH_4_-oxidizing bacteria and significant knowledge gaps related thereto.

Clustered regularly interspaced short palindromic repeats (CRISPR) interference (CRISPRi) is a genetic tool that has been leveraged for functional genetic screens in diverse bacteria^31–39^. CRISPRi uses the nuclease-deficient “dead” CRISPR-associated (Cas) variant dCas9 and a single-guide RNA (sgRNA) for the sequence-specific repression of gene expression^40^. This technology can be used to target single genes or scaled for genome-wide screening via massively parallel oligonucleotide synthesis of pooled sgRNAs. Functional genomic screens using CRISPRi are coupled with next generation sequencing to quantify the depletion or enrichment of sgRNAs (sgRNA differential fitness) during selective bacterial cultivation^41^. Notably, transcription repression by CRISPRi can be tightly regulated using inducible promoters to drive dCas9 and/or sgRNA expression; thus, in contrast to transposon mutagenesis, CRISPRi can be applied to conditionally evaluate the effect of repressing essential gene expression on bacterial growth. Recent developments of methanotroph CRISPR genome editing tools indicate that Cas9 and sgRNAs are functional in Gammaproteobacterial and Alphaproteobacterial methanotrophs^31,42–46^ paving the way for CRISPRi and forward genetic approaches in methanotrophs that have eluded these bacteria for decades.

Here, we modified a previously developed methanotroph CRISPR-Cas9 genome editing system^44^ to facilitate CRISPRi and demonstrate its utility for repressing gene transcription in the phylogenetically diverse methanotrophs *Methylococcus capsulatus* Bath and *Methylosinus trichosporium* OB3b. We performed a genome-scale CRISPRi screen using 45,798 unique sgRNAs targeting all annotated genes in the *M. capsulatus* Bath genome to investigate gene-function relationships during CH_4_ cultivation. We identified 233 genes whose transcription repression resulted in significantly decreased fitness and repression of 13 genes associated with a fitness advantage. sgRNAs with decreased fitness scores targeted genes encoding proteins involved in critical cellular processes like ribosome biosynthesis, translation, transcription, and central biosynthetic metabolism, underscoring the utility of CRISPRi for the identification of predicted and novel essential genes. The CRISPRi system and sgRNA library developed here can be used for facile gene-function analyses and genomic screening to identify methanotroph genetic determinants under an array of screening conditions. Further, these CRISPRi screening methodologies can also be applied to high-throughput engineering approaches for isolation of improved methanotroph biocatalysts.

## Materials and Methods

### Bacterial strains and cultivation

Bacterial strains used in this study are shown in Table 1. *E. coli* strains were cultured in lysogeny broth (Lennox) with 25 μg/mL kanamycin, 50 μg/mL spectinomycin, or 10 μg/mL gentamicin for transformant selection. *Methylococcus capsulatus* Bath and *Methylosinus trichosporium* OB3b frozen stocks were streaked onto nitrate mineral salts (NMS) solid medium and maintained in stainless steel gas chambers supplied with 20% CH_4_ in the gas phase at 37°C or 30°C, respectively, as previously described^44^. Methanotroph strains were passaged weekly for 4 weeks (maximum passage = 5) after which time a new culture from frozen stock was initiated. Plasmids were transferred to methanotrophs via biparental mating by spreading equivalent biomass of S17-1λ *E. coli* and recipient methanotroph biomass on NMS mating agar mating plates and incubating in a 20% CH_4_ atmosphere for 24 h as previously described ^44^. After mating, methanotroph transformants harboring plasmids were selected on NMS medium containing 50 μg/mL spectinomycin (pJHCi and derivatives), 10 μg/mL gentamicin (pDS1), and/or 50 μg/mL kanamycin (pJHint). Methanotrophs were cultured in 150 mL vials containing 10 mL of NMS medium at 37°C (*M. capsulatus*) or 30°C (*M. trichosporium*) at 200 rpm orbital shaking. After inoculation with plate-derived biomass to OD_600_ 0.01, vials were crimped with gray butyl stoppers to create gas-tight seals followed by CH_4_ addition to the headspace via syringe to reach a final CH_4_ concentration of 20% in air (v/v). Where applicable, sgRNA and/or dCas9 expression were induced with 0.2 µg/mL anhydrotetracycline (aTc). High resolution bacterial growth was measured every 20 seconds by Cell Growth Quantifier (CGQ) optical sensors (Scientific Bioprocessing), and the data is presented as the mean arbitrary light backscatter units as directly detected by the sensors.

**Table 1.**
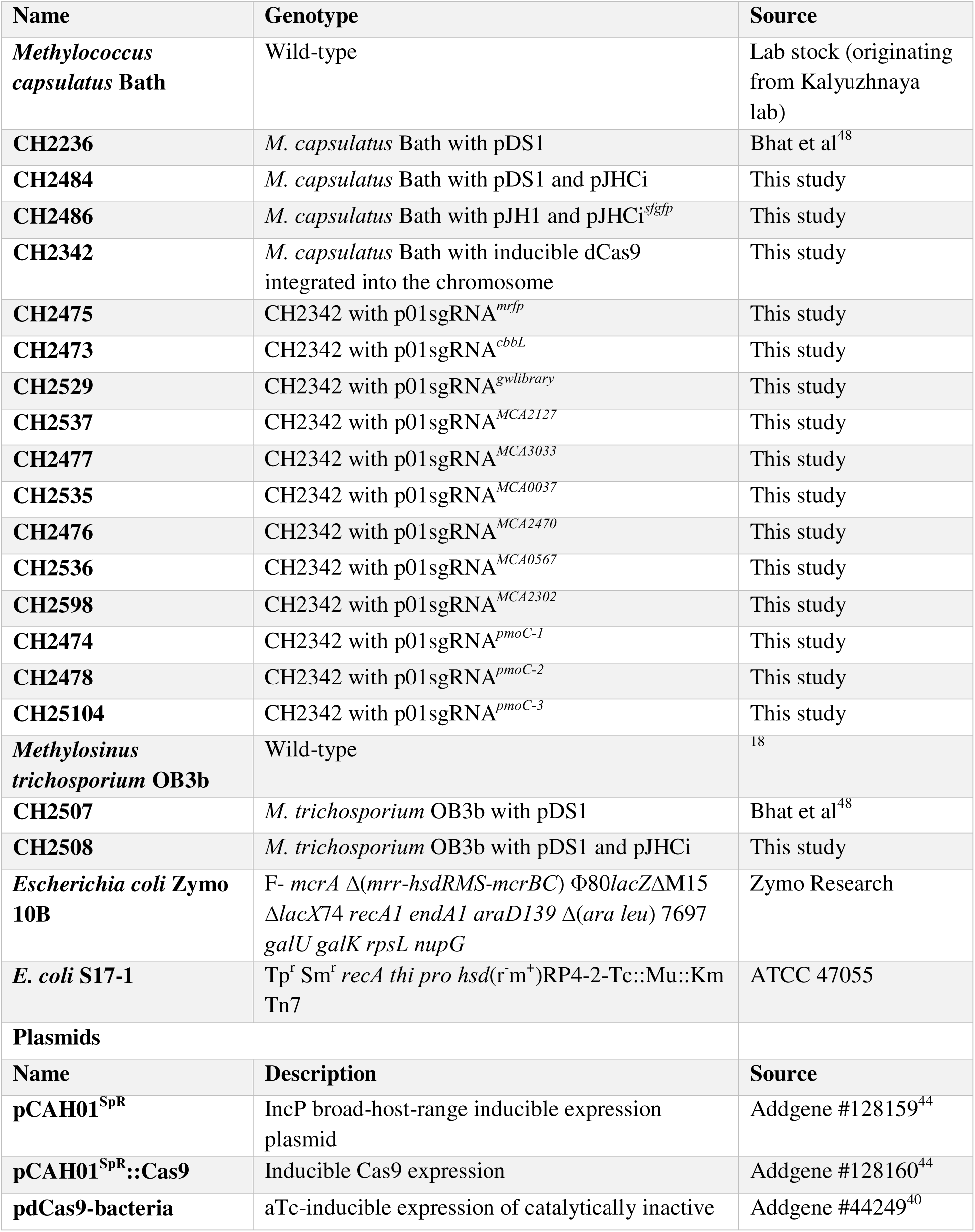
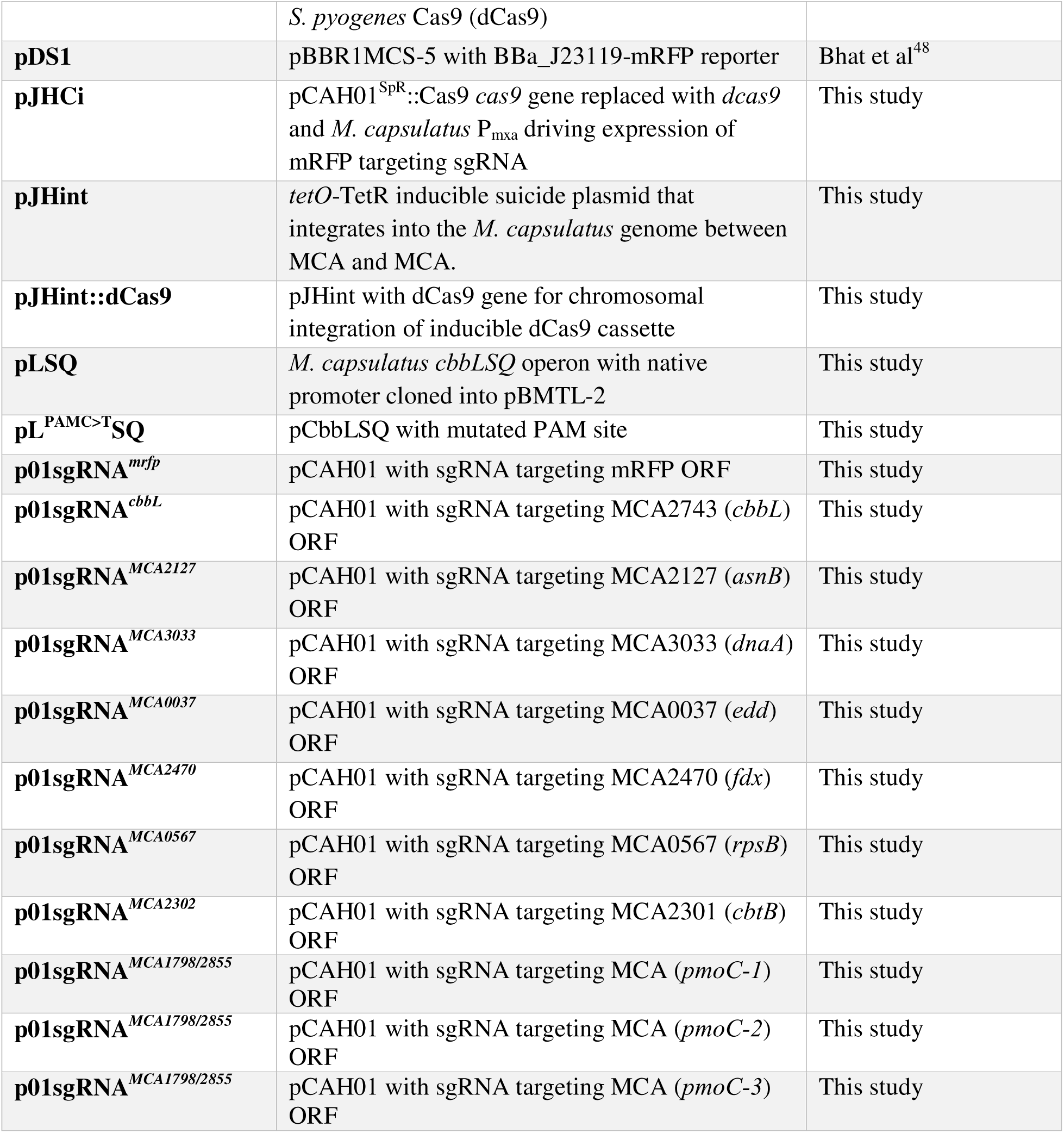
Strains and Plasmids.

### CRISPRi plasmid construction and sgRNA exchange

Primers and other synthetic DNA used in this study are shown in Table 2. To construct a broad-host-range, single plasmid CRISPRi system, we amplified a P*_mxaF_*-sgRNA with an mRFP spacer sequence from the pgRNA plasmid^44^ using primers oCAH267/289 and ligated it with pCAH01^SpR^::Cas9^44^ linearized with primers oCAH230/231 via isothermal assembly using HiFi Gibson Assembly Master Mix (New England Biolabs). We then amplified the dCas9 gene from plasmid pdCas9-bacteria^40^ with oCAH952/1507 and assembled it with the above plasmid backbone amplified with oCAH4/3, exchanging the Cas9 gene with dCas9 to generate plasmid pJHCi. The aTc-inducible suicide plasmid pJHint was constructed via isothermal assembly of the following six DNA fragments: 1) pUC origin of replication from pCAH01 amplified with oCAH403/404; 2) 1kb upstream homology arm amplified with oCAH405/406 from *M. capsulatus* genomic DNA; 3) P*_tet_* promoter/operator amplified from pCAH01 with oCAH407/408 ; 4) *tetR*-kn^R^ cassette from pCAH01 amplified with oCAH409/410; 5) 1kb downstream homology arm amplified with oCAH411/412 from *M. capsulatus* genomic DNA; and 6) *oriT* origin of transfer amplified from pCAH01 with oCAH413/414. pJHint was then amplified with primers oCAH3/4 and assembled with the dCas9 gene fragment to generate plasmid pJHintdCas9. pJHintCas9 was transferred to *M. capsulatus* Bath via conjugation and transformants with the P*_tet_*-dCas9 module integrated into an intergenic site between MCA_tRNA-Ile and MCA_tRNA-Pro were selected on NMS solid medium containing 25 µg/mL kanamycin. Successful integration was confirmed via PCR using primers oCAH447/448 that flank the integration site. For sgRNA expression, we initially amplified the mRFP targeting sgRNA from pJHCi with primers oCAH1083/1081 and cloned it into pCAH01 amplified without a ribosomal binding site with primers oCAH4/119 to generate p01sgRNA*^mrfp^*. Other aTc-inducible sgRNA expression plasmids were generated by exchanging the spacer sequence via isothermal assembly of a single-stranded oligo containing a unique spacer and the pCAH01sgRNA*^mrfp^* amplified with primers oCAH1190/119.

**Table 2.**
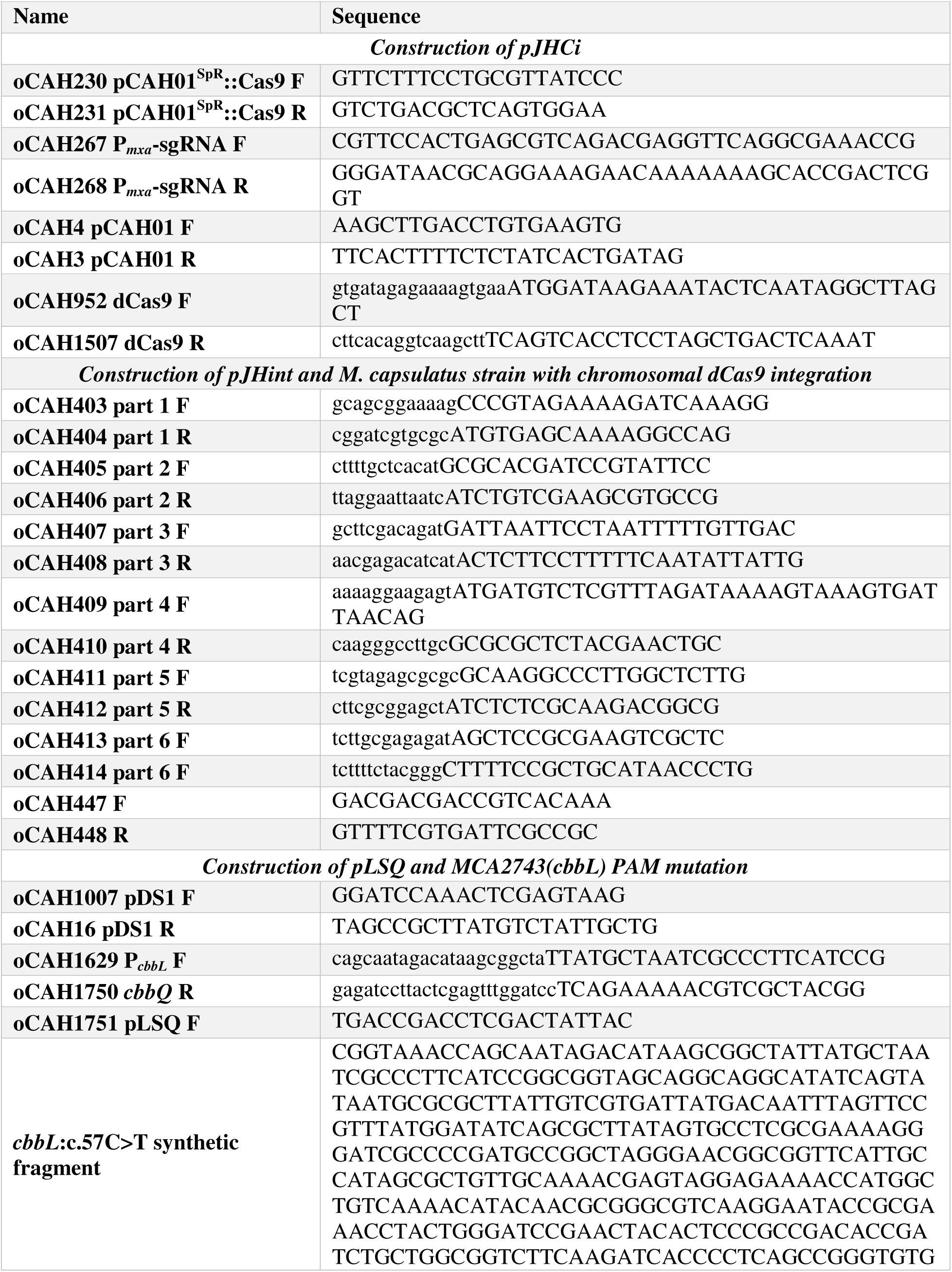
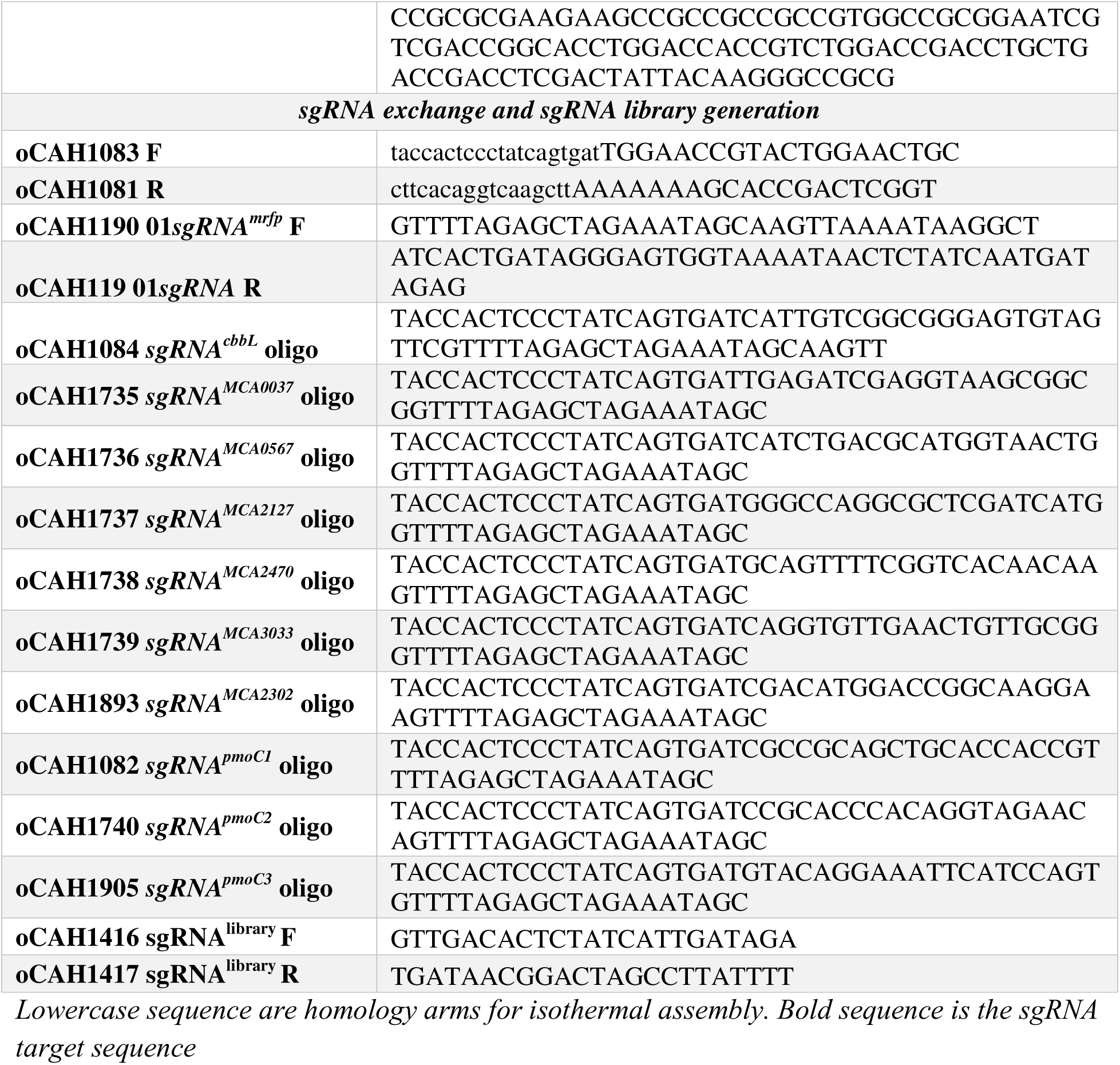
Primers and synthetic DNA fragments.

### Fluorescence measurements

pJHCi was transferred to *M. capsulatus* Bath or *M. trichosporium* OB3b harboring the pDS1 mRFP reporter plasmid via conjugation. Alternatively, pDS1 and p01sgRNA*^mrfp^* were iteratively transferred to the *M. capsulatus* Bath strain with chromosomally integrated P*_tet_*-dCas9. Overnight seed cultures of these strains were subcultured to OD_600_ = 0.01 in liquid medium containing appropriate antibiotics with or without aTc induction and cultivated for 72h to ∼ OD_600_ 2.0. 200 µL culture was transferred to a 96-well microplate, and mRFP1 fluorescence (ex_532nm_, em_588nm_, gain = 80) and optical density (A_600nm_) was measured with a BioTek Synergy Mx microplate reader. The data were normalized by dividing raw fluorescence reads by the optical density. mRFP expression was also monitored via fluorescence microscopy by transferring 5 µL *M. capsulatus* logarithmic growth phase cultures with or without CRISPRi induction to a glass slide and mounted with a cover slip. RFP (ex575-640nm, em629nm) images were acquired using a Zeiss AXIO Imager M2 fluorescence microscope with 100X (oil) Plan-Apochromat (HC DIC) objective and processed with ZEN software.

### Construction of pLSQ and cbbL PAM mutant

The *M. capsulatus* Bath *cbbLSQ* operon (MCA2743-2746) including 150bp upstream of *cbbL* presumed to contain the *cbbL* promoter was amplified from purified genomic DNA using primers oCAH1629 and oCAH1750. The amplicon was assembled with pDS1 amplified with primers oCAH1007 and oCAH16 to generate pLSQ. This plasmid was then linearized with primers oCAH1751 and oCAH16 and assembled with a synthetic fragment containing a *cbbL*:c.57C>T protospacer adjacent motif (PAM) mutation to generate pLSQ^ΔPAM^. The pCAH01sgRNA*^cbbL^* plasmid was transferred *to M. capsulatus* Bath harboring the pLSQ or pLSQ^ΔPAM^ strains and growth was assessed with or without aTc induction using CGQ optical sensors.

### Design, synthesis, and construction of the M. capsulatus genome-wide sgRNA library

Synbio Technologies designed and synthesized the sgRNA oligo pool following design rules to identify 15 PAM and target sequences per the predicted 3,034 open reading frames in the NC_002977.6 RefSeq genome and to preferentially target the non-template DNA strand within the first 50% of the 5’ end of the target gene. The sgRNA library consisted of 45,798 sgRNA oligos (120 bp) with each containing a unique spacer sequence and 5’ and 3’homologous sequence for facile cloning into the p01sgRNA plasmid. The library also included 400 non-targeting sgRNAs with randomly generated spacers. The pooled sgRNA library was amplified via PCR using primers oCAH1416/1417 and Q5 polymerase (New England Biolabs) following the manufacturers recommended reaction conditions for 20 cycles to limit biased amplification. The amplicons were assembled with p01sgRNA*^mrfp^* amplified with oCAH1190/119 via isothermal assembly and transformed into high-efficiency DH10b competent cells (New England Biolabs). Prior to plating on selective medium, the competent cells were serially diluted to determine transformant colony forming units (CFU/mL). 1.02 x 10^7^ DH10b transformants were obtained after transformation, three orders of magnitude greater than the synthesized library size. Twenty antibiotic-resistant transformants were screened via PCR and Sanger sequencing to determine p01sgRNA*^mrfp^*background, which was < 5 %. The transformants were then pooled and the library miniprepped using the Monarch Plasmid Miniprep Kit (New England Biolabs). The mixed p01sgRNA library was then transferred to chemically competent S17 *E. coli* obtaining 1.15 x 10^6^ S17 transformants, which were conjugated with *M. capsulatus* Bath strain with integrated P*_tet_*-dCas9 to generate the methanotroph CRISPRi library. *M. capsulatus* Bath transformants were quantified via serial dilution plating and pooled for genomic DNA extraction and long-term - 80°C storage. 6.8 x 10^7^ transconjugants were obtained, 1000X the amount of unique sgRNAs in the synthesized library. Library composition was determined via Oxford Nanopore Technologies amplicon sequencing as described below.

### Oxford Nanopore Technologies sequencing and bioinformatic analysis

The sgRNA library was cultivated in liquid NMS with or without CRISPRi induction. After 72 h of selection, genomic DNA was extracted from an OD = 2 equivalent cell pellet using the Zymo DNA extraction kit following the manufacturers protocol. 50 ng pooled sgRNA library was used as a template to amplify a ∼250 bp amplicon containing the sgRNA sequence with Q5 polymerase using primers oCAH1416/1417 and the same reaction conditions as above (20 cycles). The amplicons were barcoded with the native barcoding kit 24 V14 following the manufacturers protocol (Oxford Nanopore Technologies, ONT) and sequenced using a MinION sequencing device with high accuracy basecalling. Fastq sequences were then trimmed and filtered using Geneious Prime software. The 20 bp sgRNA target sequences were extracted from the trimmed and filtered sequence reads using a custom Python script to process ONT sequence data stored as compressed .fasta.gz files. The script performs a two-step search for the constant sgRNA sequences (TATTTTACCACTCCCTATCAGTGAT, TTTTAGAGCTAGAAATAGCAAGTT or their reverse complements ATCACTGATAGGGAGTGGTAAAATA, AACTTGCTATTTCTAGCTCTAAAAC) flanking the target sequence and extracts the adjacent 5’ or 3’ 20 bp target sequence from the read dependent on the identified query sequence/read strand orientation. The script utilizes pairwise alignment to refine the search results, ensuring accurate extraction of relevant sequences. After extraction, the sgRNA target sequences were aligned to either a custom reference library containing the 45,798 sgRNA library sequences or the *M. capsulatus* Bath genome (NC_002977.6); mapped reads were quantified and normalized as sgRNA target sequence per million (TPM); and the fitness score (sgRNA target sequence differential abundance between induced and uninduced samples) was determined following a DEseq2 workflow using Geneious software. Enrichment analysis of the most differentially abundant sgRNA targets was performed using ShinyGo 0.82 using the STRING database^47^.

### Co^2+^-deplete cultivation, gas chromatography and biomass productivity calculations

Wild- type *M. capsulatus* solid medium-derived biomass was inoculated into acid-washed serum vials containing 10 mL NMS medium with decreasing CoCl_2_ in the trace elements to OD_600_ = 0.01 and growth was measured with CGQ optical sensors. All medium components were made with HPLC-grade water in acid-washed glassware. The CH_4_ in the headspace was determined 10 min after gas addition and every 24 h during methanotroph cultivation using an SRI 8610c gas chromatograph with TCD and FID detectors (SRI Instruments). CH_4_ was quantified as % atmospheric composition by comparison to known standards and converted to weight using the known headspace volume and specific gravity (0.622 mg/mL at 37°C). Dry cell weight (DCW) was determined by converting OD_600_ to DCW using a formula empirically determined by comparing OD_600_ measurements to colony forming units during serum vial cultivation (OD_600_ = 1 is equivalent to 0.254 ± 0.0249 g DCW/L).

### Illustrations

Data were graphed using GraphPad Prism 10 software. Circos plots were generated using Circa (https://circa.omgenomics.com/). Workflow schematics were created with BioRender.com.

### Statistical analysis

Statistical analysis was performed using GraphPad Prism 10 software. The data between two groups were analyzed using unpaired t tests. Data were considered statistically significant when *p* ≤ 0.05.

## Results and Discussion

### Construction and validation of methanotroph CRISPRi tools

We previously developed a two-plasmid, broad-host-range CRISPR-Cas9 system that enabled *M. capsulatus* genome editing ^44^. This two-plasmid system consisted of a 1) pBBR-based plasmid with the *M. capsulatus* constitutive PQQ-dependent methanol dehydrogenase P*_mxaF_*promoter driving single guide RNA (sgRNA) expression, and a compatible 2) IncP-based plasmid, pCAH01, with anhydrotetracycline-inducible Cas9 expression (pCAH01^SpR^::*cas9*)^44^. To construct a single plasmid CRISPRi system, we moved the P*_mxaF_*-sgRNA module to pCAH01^SpR^::*cas9* and replaced the *cas9* gene with the dead cas9 (*dcas9*) gene encoding the nuclease-deficient dCas9 variant (D10A, H840A) creating pJHCi (Figure 1a). The ability of the CRISPRi system to repress transcription in the Gammaproteobacterial methanotroph *M. capsulatus* Bath and Alphaproteobacterial methanotroph *Methylosinus trichosporium* OB3b was tested using strains expressing an mRFP reporter (Figure 1b)^48^. pJHCi with a sgRNA targeting the non-template strand of the mRFP open reading frame (sgRNA^mRFP^) was transferred to the methanotroph reporter strains via conjugation. The CRISPRi system reduced *M. capsulatus* and *M. trichosporium* mRFP fluorescence by 6.3-fold and 2.3-fold, respectively, after 72 h of bacterial cultivation in liquid NMS with CH_4_ as the carbon source (Figure 1c). The decreased efficiency of the CRISPRi system in *M. trichosporium* may be related to differences in sgRNA and/or dCas9 abundance as promoters derived from *Gammaproteobacteria*, like those driving transcription of the sgRNA (*M. capsulatus* P*_mxa_*) and dCas9 (P*_tet_*), exhibit lower activity in *M. trichosporium* compared to *M. capsulatus*^48^.

**Figure 1.**
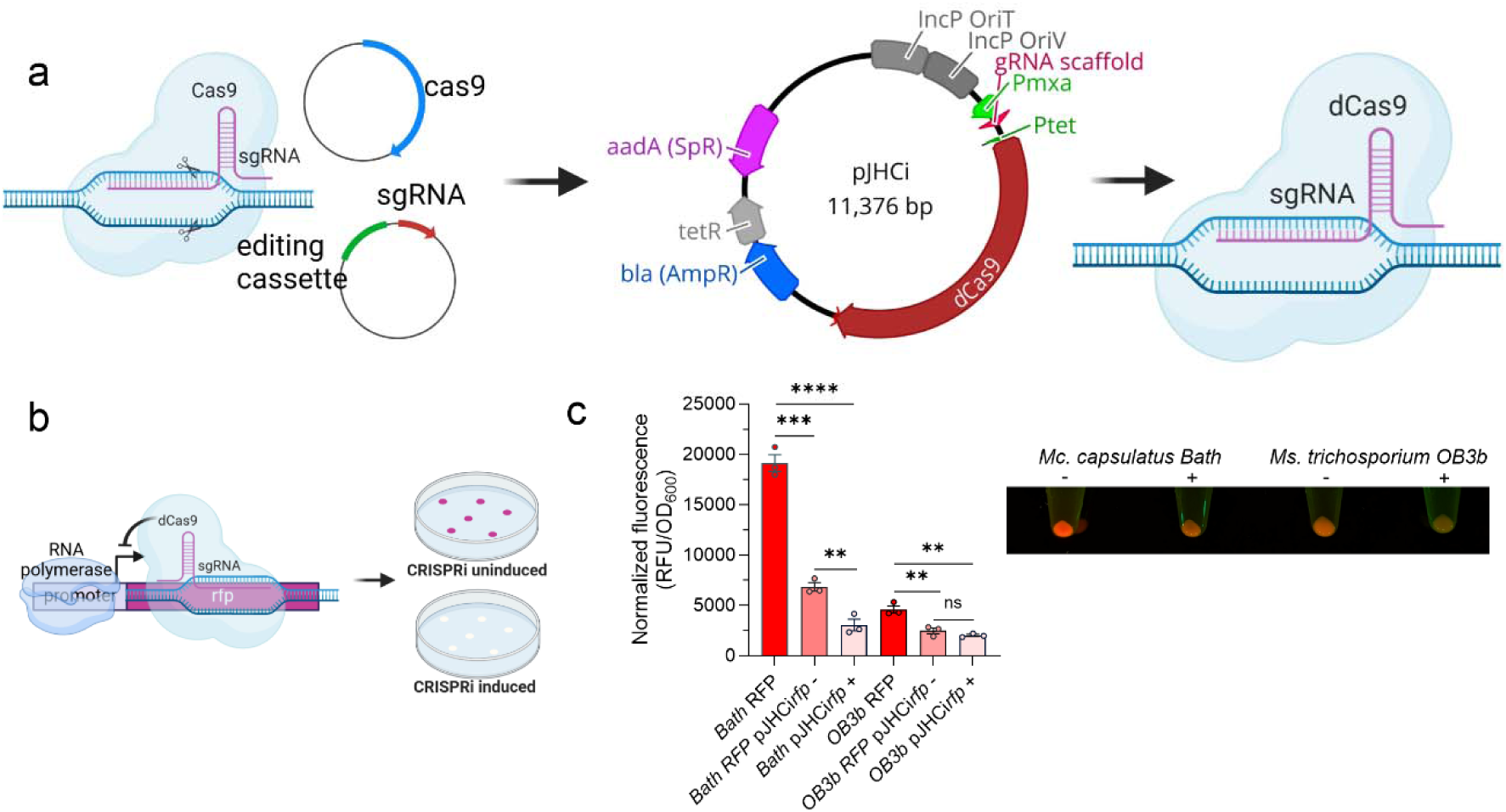
Construction and validation of a methanotroph CRISPRi system. a) Schematic overview of converting a two-plasmid CRISPR genome editing system into a single-plasmid CRISPRi system for targeted repression of gene expression. b and c) The mRFP fluorescent reporter was used as a read-out of CRISPRi functionality in Gammaproteobacterial methanotroph *Methylococcus capsulatus* Bath and Alphaproteobacterial methanotroph *Methylosinus trichosporium* OB3b with (+) or without (-) dCas9 induction by anhydrotetracycline (aTc). A representative image of cell pellets is shown. Data in c represent the mean ± SEM from two independent experiments (n=3-5). ** *p* ≤ 0.01, *** *p* ≤ 0.001, **** *p* ≤ 0.0001.

The pJHCi CRISPRi system limited the production of fluorescent protein in the absence of dCas9 induction in both methanotrophs (Figure 1c), suggesting that low dCas9 expression from leaky P*_tet_* promoter gene transcription in the context of high, constitutive sgRNA expression is sufficient for transcription repression. The leaky functionality of the CRISPRi system is not desirable for functional genomic applications where sgRNA expression could cause fitness defects during strain or library generation. Thus, to develop a system with more tightly regulated CRISPRi repression, we designed a configuration wherein both dCas9 and the sgRNA are inducible (Figure 2a). mRFP fluorescence in this strain was similar to a control strain with the mRFP reporter plasmid alone (Figure 2b), supporting that the CRISPRi system did not inhibit mRFP transcription in the absence of aTc induction. In contrast, simultaneous induction of dCas9 and sgRNA^mRFP^ led to a 3.3-fold decrease in mRFP fluorescence (Figure 2b).

**Figure 2.**
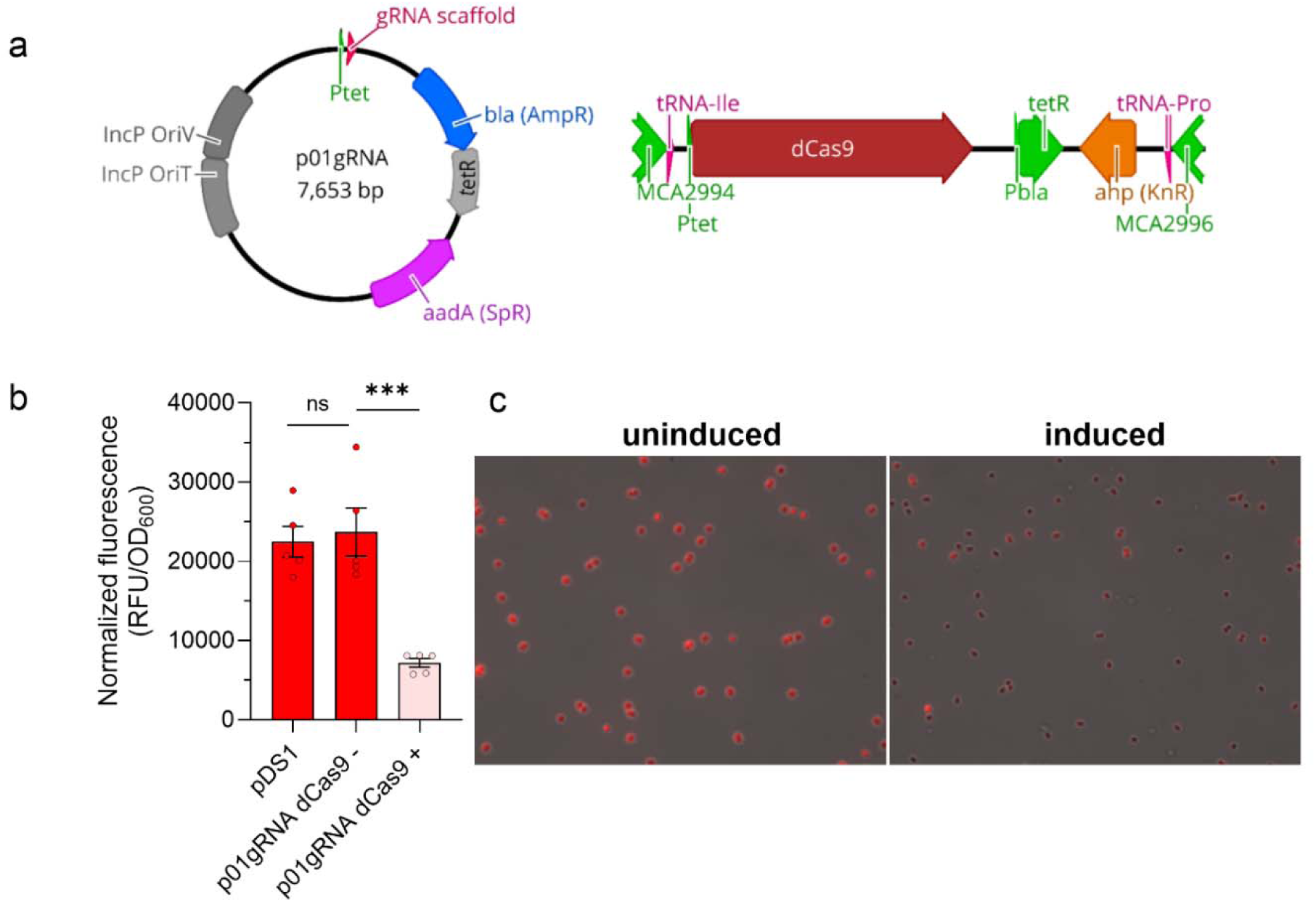
A CRISPRi system with inducible dCas9 and sgRNA enables fine-tuned transcription repression in *M. capsulatus*. a) The pJHCi single plasmid system exhibited leaky functionality, so an *M. capsulatus* Bath strain was generated with an aTc-inducible dCas9 cassette integrated into the chromosome and plasmid with aTc-inducible sgRNA expression. b) mRFP fluorescence with or without aTc induction compared to the pDS1 mRFP reporter plasmid control strain showing tightly regulated CRISPRi gene transcription repression. c) Fluorescence microscopy of *M. capsulatus* with or without CRISPRi induction for mRFP repression. Representative images from biological replicates are shown. Data in b represent the mean ± SEM from two independent experiments (n=3-5). ns-not significant, *** *p* ≤ 0.001.

Fluorescence microscopy was used to assess CRISPRi efficiency in individual cells. As shown in Figure 2c, CRISPRi repressed mRFP fluorescence in most of the bacterial population, but some cells exhibited high mRFP fluorescence indicating a small percentage of the population may “escape” CRISPRi repression.

### CRISPRi-mediated transcription repression of *M. capsulatus* RubisCO causes growth inhibition

We previously showed that the RubisCO enzyme encoded by *M. capsulatus* is essential for its growth on CH ^49^. As such, we targeted the *cbbL* gene encoding the RubisCO large subunit to test the ability of the CRISPRi system to repress transcription from the *M. capsulatus* chromosome. Transconjugants could only be obtained in the absence of sgRNA*^cbbL^*and dCas9 induction (Figure 3a), supporting the essentiality of RubisCO for *M. capsulatu*s growth^49^. Transconjugants isolated in the absence of induction were subsequently cultivated in liquid NMS with or without the aTc inducer, and high-resolution growth analysis showed that CRISPRi-repression of *cbbL* transcription significantly inhibited bacterial growth (Figure 3b). Notably, the induced culture began to expand after five days and exhibited typical logarithmic growth kinetics thereafter (Figure 3b). The aTc inducer has a 17 h half-life in minimal medium at pH 7 and 37°C ^50^, so we hypothesized that the delayed growth observed in induced samples may be due to ineffective aTc concentrations after five days cultivation at 37°C. However, we did not observe a growth defect when the induced, outgrown sgRNA*^cbbL^* culture was diluted into fresh induction medium (data not shown), indicating that the CRISPRi system is no longer effective in this expanded population. Known mutations in the dCas9 gene^51^ or others in the sgRNA, P*tet-tetO* promoter, or the *tetR* gene that would render the CRISPRi system nonfunctional were not identified in five “escapee” colonies via Sanger sequencing (data not shown), so the underlying mechanism of resistance to CRISPRi repression in these strains remains elusive. Given the essentiality of RubisCO for *M. capsulatus* growth, it is likely there is a strong selective pressure to overcome *cbbL* transcription repression in these cells.

**Figure 3.**
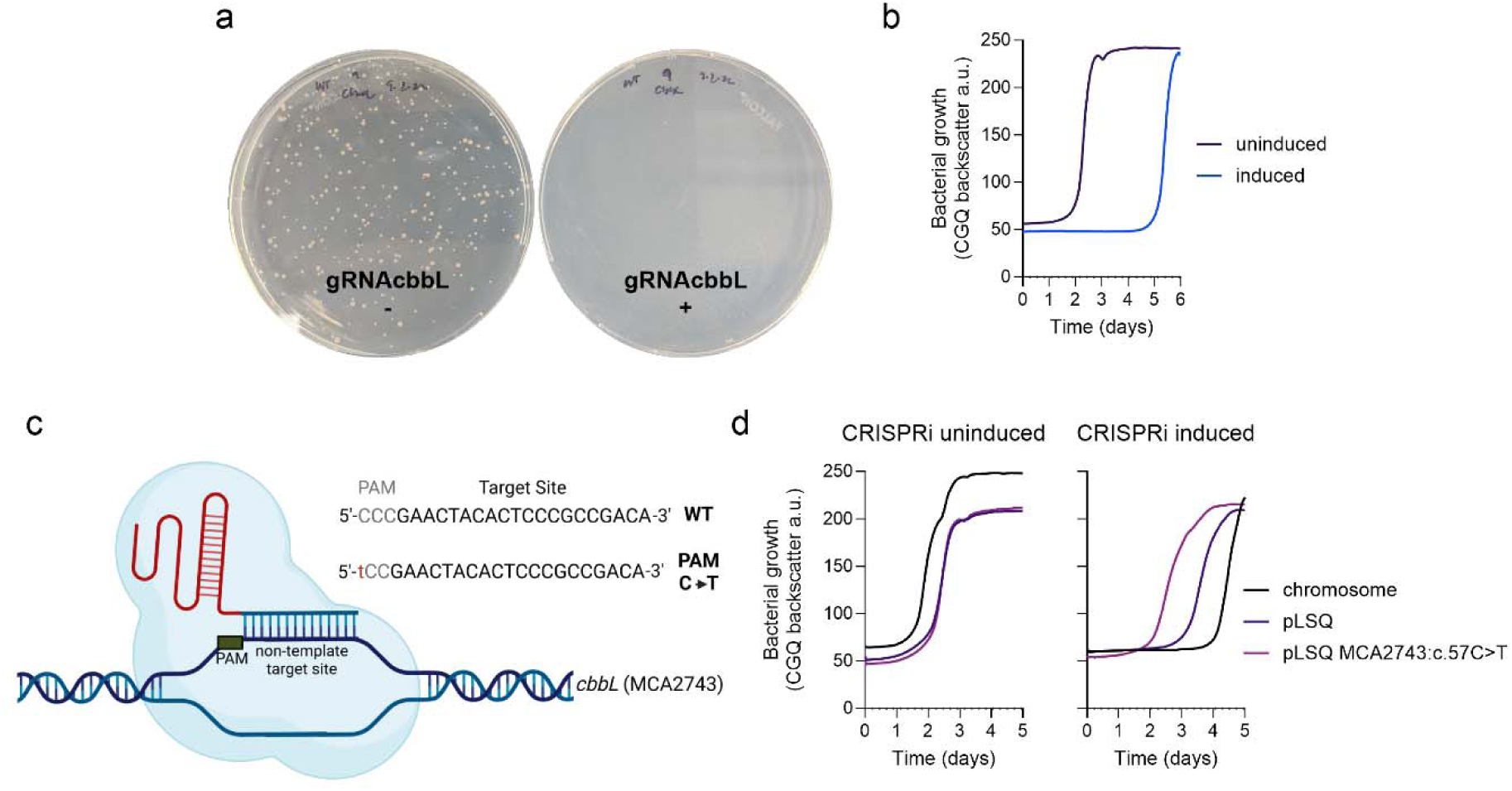
CRISPRi-mediated transcription repression of the genes encoding the essential ribulose 1,5 bisphosphate carboxylase/oxygenase (RubisCO) enzyme limit *M. capsulatus* growth. Growth of *M. capsulatus* with (+) or without (-) induction of dCas9 and an sgRNA targeting the *cbbL* gene encoding the large subunit of RubisCO on solid NMS medium (a) or in liquid NMS medium in a serum vial over time (b) with a 20% CH_4_ in air atmosphere. c) A complementing plasmid expressing the *cbbLSQ* operon (pLSQ) was constructed and the protospacer adjacent motif (PAM) site in the *cbbL* target sequence (MCA2743) was mutated (c57C>T). d) Growth of strains carrying the pLSQ or pLSQ^ΔPAM^ compared to an isogenic background control strain with or without CRISPRi induction. Data in b and d represent the means of two independent high-resolution cultivation experiments (n=6).

To further test the specificity of the CRISPRi system, we constructed a plasmid containing the *cbbLSQ* operon under control of its native promoter region (pLSQ) and another plasmid with a mutated *cbbL* target protospacer adjacent motif (PAM) (MCA2743:c.57C>T, pLSQ^ΔPAM^) expected to prevent dCas9 DNA binding^5253^ (Figure 3c). CRISPRi targeting of *cbbL* in the pLSQ^ΔPAM^ strain did not inhibit growth (Figure 3d), supporting that expression of RubisCO from this plasmid was not inhibited and able to complement chromosomal *cbbLSQ* transcription repression. In contrast, CRISPRi-mediated repression of *cbbL* transcription in the strain with pLSQ inhibited bacterial growth, but not to the extent of that observed in the strain lacking the plasmid (Figure 3d). The pLSQ plasmid has a pBBR-based replicon, which is maintained in *M. capsulatus* at ∼50 copies/cell^48^. As such, the difference in growth inhibition observed between the pSLQ and control strain may be correlated to an increase in target sites in cells harboring pSLQ, thereby limiting CRISPRi transcription repression efficiency compared to targeting a single chromosomal site.

### CRISPRi functional genomic screening identifies genes essential for CH_4_-dependent growth

To evaluate the utility of our CRISPRi system for functional genomic screening, we designed and synthesized a pooled sgRNA library consisting of 45,798 sgRNAs, ∼15 sgRNAs per *M. capsulatus* protein-coding genes (3,034 annotated in the NC_002977.6 RefSeq genome) and 400 non-targeting control sgRNAs (supplemental Table S1-S2). Eighteen open-reading frames had less than 15 sgRNAs in the library and zero sgRNA target sequences were identified for 3 genes (MCA0050, MCA0307, MCA1517, supplemental Table S3). The workflow for cloning and screening the sgRNA library is shown in Figure 4a. We determined the *M. capsulatus* library coverage by amplifying the sgRNAs from genomic DNA isolated from 6.8 x 10^7^ pooled transconjugants. The sgRNA amplicons were sequenced using an Oxford Nanopore Technologies MinION device; 680,425 reads with QC 12+ scores were trimmed and filtered using Geneious software followed by extraction of the 20 bp sgRNA targeting sequences from the processed sequences using a custom script. In total, 680,304 extracted sgRNA spacers were aligned to the synthesized genome-wide CRISPRi sgRNA library reference with 45,554 of the 45,798 synthesized sgRNAs detected (99.5% sgRNA library coverage, supplemental Table S4) with at least one sgRNA per predicted DNA coding sequence (except the three genes originally not included in the synthesized library, supplemental Table S5).

**Figure 4.**
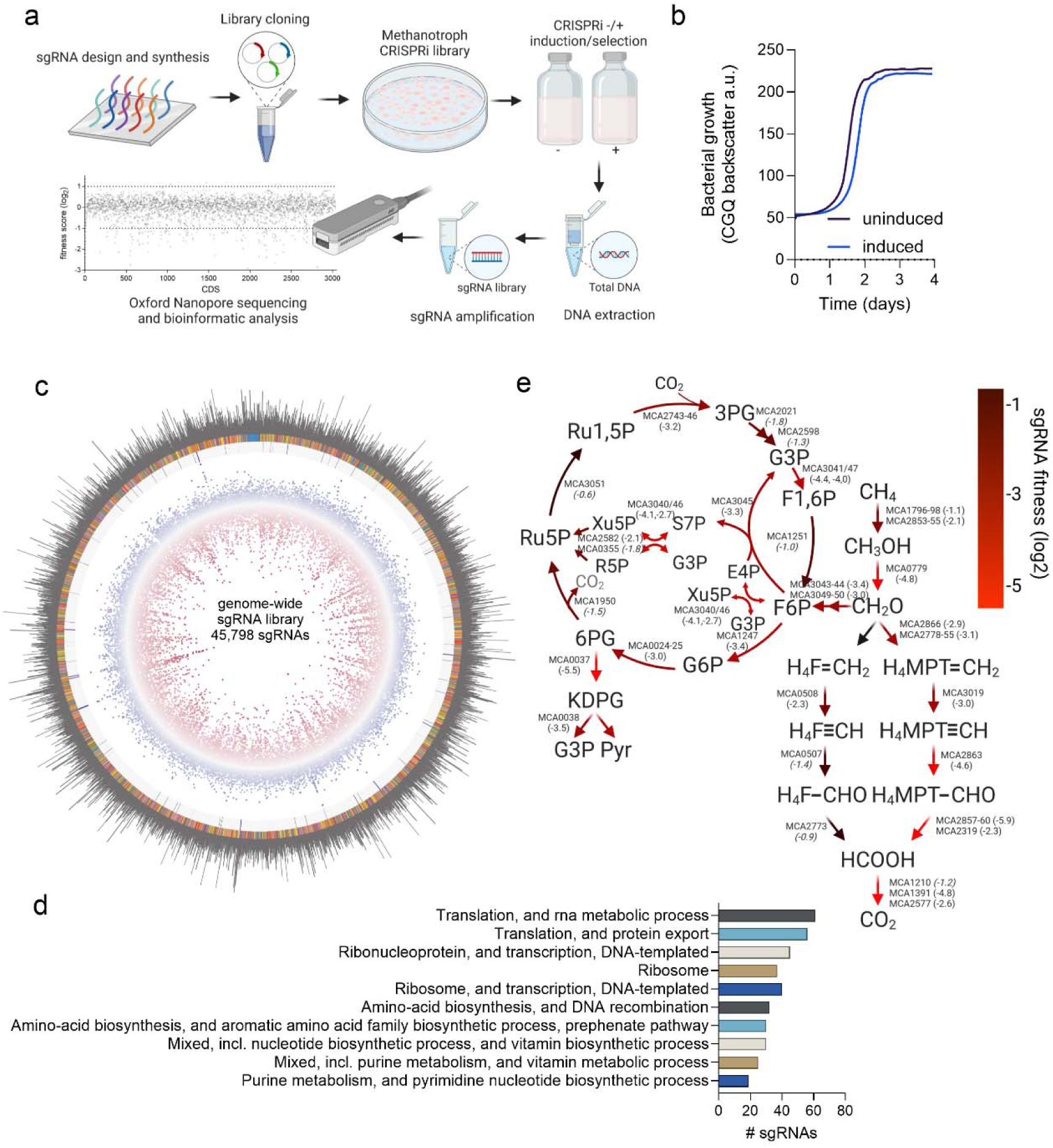
A CRISPRi genome-wide screen identifies genes essential for CH_4_-dependent growth of *M. capsulatus*. a) 45,798 sgRNAs were synthesized via massively parallel oligonucleotide synthesis and cloned into the inducible plasmid pCAH01 to generate a cloned sgRNA library. The sgRNA library was transferred to an *M. capsulatus* Bath strain with a chromosomally integrated P*_tet_*-dCas9 cassette via biparental mating. b) The library was cultivated with or without sgRNA and dCas9 induction in NMS medium in serum vials with a 20% CH_4_ in air atmosphere. After selection, the sgRNA composition and abundances were determined via Oxford Nanopore Technologies amplicon sequencing and bioinformatic analysis to quantify sgRNA fitness. c) Circos plot of the methanotroph sgRNA library raw reads (outer ring) mapped to the 45,798 synthesized sgRNA oligos (rainbow ring with non-targeting sgRNAs in blue). Differential sgRNA fitness scores determined via DeSeq2 between CRISPRi induced (n=4) and uninduced (n=4) samples during selection/cultivation with CH_4_ as the sole carbon and energy source (4^th^ ring, blue dots log2 >1; red dots log2 <1). sgRNAs depleted in the screen (negative fitness scores) correspond to genes with transcription repression causing a growth defect where those enriched (positive fitness scores) correspond to genes with transcription repression causing a growth advantage. sgRNAs with statistically significant (*p* ≤ 0.01) differential fitness scores are indicated (blue squares, 3^rd^ ring). d) ShinyGo enrichment analysis of the 429 genes with sgRNA log2 fitness scores ≤ -3 and *p* ≤ 0.001. e) CH_4_ assimilatory and dissimilatory metabolic pathways predicted by sgRNA fitness scores. The mean sgRNA fitness scores are indicated in parentheses next to the gene loci that encode metabolic enzymes. 3PG, 3-phosphoglycerate; G3P, glyceraldehyde 3-phosphate; F1,6P, fructose 1,6-bisphosphate; F6P, fructose 6-phosphate; E4P, erythrose 4-phosphate; Xu5P, xylulose 5-phosphate; S7P, sedoheptulose 7-phosphate; R5P, ribose 5-phosphate; Ru5P, ribulose 5-phosphate; R1,5P, ribulose 1,5-bisphosphate; G6P, glucose-6-phosphate; 6PG, 6-phosphogluconate; KDPG, 2-keto-3-deoxy-6-phosphogluconate; Pyr, pyruvate; H_4_F=CH_2_, methylene-tetrahydrofolate; H_4_MPT=CH_2_, methylene-tetrahydromethanopterin; H_4_F≡CH, methenyl-tetrahydrofolate; H_4_MPT≡CH, methenyl-tetrahydromethanopterin; H_4_F-CHO, formyl-tetrahydrofolate; H_4_MPT-CHO, formyl-tetrahydromethanopterin. Scores with *p* ≥ 0.01 are italicized. Data in b represent the means of two independent high-resolution cultivation experiments (n=6).

The methanotroph genome-wide sgRNA library was cultivated in serum vials with an atmosphere of 20% CH_4_ in air with and without CRISPRi induction to identify genes essential for *M. capsulatus* growth. We observed a minor growth defect of CRISPRi induced cultures compared to uninduced cultures, but logarithmic growth kinetics and entry into stationary phase were similar between the conditions (Figure 4b). The sgRNA library compositions in CRISPRi induced and noninduced samples 72 h after selection were quantified by sgRNA amplicon sequencing and mapping to both the synthesized sgRNA library and the *M. capsulatus* genome (supplemental Tables S6-S13, read counts). We identified between 61.1-91.8% of the synthesized sgRNAs in the sample sequences, with increased coverage strongly correlated to sequencing depth (supplemental Figure S1, R^2^ = 0.93). Although complete library coverage was not obtained in the screening samples with the sequencing depth achieved here, all gene targets were represented in the sequence data with 3,027/3,034 (99.8%) coding DNA sequences with ≥10 guides represented (supplemental Tables S6-S13). Comparing the sgRNA library composition between CRISPRi induced vs noninduced samples showed that 1,019 sgRNAs were depleted (log2 fitness score ≤ 1, *p* ≤ 0.001) and 59 sgRNAs were enriched (log2 fitness score ≤ 1, *p* ≤ 0.001) after 72 h of selection, presumably corresponding to a fitness defect or advantage phenotype associated with transcription repression of the sgRNA target gene (Figure 4c and Table S14). Enrichment analysis of 429 genes with sgRNA log2 fitness scores ≤ -3 and probability values ≤ 0.001 indicated that these genes encode proteins associated with essential cellular processes like translation, transcription, ribosome biosynthesis, and amino acid and nucleotide biosynthesis (Figure 4d and Table S15). Enrichment analysis of the genes whose transcription repression resulted in a fitness advantage did not enrich any pathways using the available ShinyGO databases. Notably, 99.5% (398/400) of the non-targeting control sgRNAs did not have significant differential fitness scores (log2 ≤-1 or ≥1, *p* ≤ 0.001) between the two conditions (supplemental Table S14).

Several sgRNAs targeting known central CH_4_ metabolic enzyme genes had significant negative fitness scores (Figure 4e and supplemental Table S14), including those targeting the particulate methane monooxygenase operons (MCA1796-98 and MCA2853-55), the calcium- and PQQ-dependent methanol dehydrogenase (MCA0779), and the RuMP cycle operons (MCA3040-45 and 3046-50). As expected based on our initial CRISPRi experiments (Figure 3), sgRNAs targeting the *cbbLSQ* operon (MCA2743-46) were among those with the lowest fitness scores. Further, the sgRNAs targeting the Entner-Doudoroff pathway *edd* and *eda* genes (MCA0037-38) were significantly depleted in the screen, consistent with biochemical data and genome-scale models predicting this pathway to be essential and the primary route for pyruvate biosynthesis in *M. capsulatus* Bath^5,54,55^. Also, the depletion of sgRNAs targeting the tetrahydromethanopterin formaldehyde dissimilatory pathway corroborate flux balance model predictions that this is the primary route used by *M. capsulatus* for formaldehyde conversion to formate^54,55^. Notably, sgRNAs targeting genes encoding serine cycle enzymes showed no differential fitness during growth on CH_4_.

Many sgRNAs targeting the same gene or genes within operons exhibited similar fitness scores, highlighting the potential of the CRISPRi library and screening pipeline to identify functional gene targets (supplemental Table S14). For example, sgRNAs targeting ribosomal protein operon genes MCA2368, MCA2371, MCA2372, and MCA2374 were consistently depleted under CRISPRi induction conditions with log2 fitness scores of -5.47, -5.57, -5.70, and -6.37, respectively, while sgRNAs targeting the cobalt transporter and cobalamin biosynthetic operon genes MCA2300, MCA2301, and MCA2302 were significantly enriched with log2 fitness scores of 3.10, 2.70 and 3.65, respectively. However, significant fitness score variances between sgRNAs targeting the same gene were observed, suggesting CRISPRi effectiveness in repressing transcription can vary depending on the sgRNA target sequence. Further evaluation of the individual *pmoCAB-* and *cbbLSQ*-targeting sgRNAs showed highly variable fitness scores, especially those targeting, *pmoC*, *pmoA*, and the *cbbL* genes (supplemental Figure S2a). To confirm this variability, we tested the ability of three *pmoC*-targeting sgRNAs with variable fitness scores (sgRNA*^pmoC^*^1^:log2 = 0.34; sgRNA*^pmoC^*^2^: log2 = -1.11; sgRNA*^pmoC^*^3^: log2 = -2.10) to inhibit *M. capsulatus* growth. The extent of bacterial growth inhibition was correlated to the respective sgRNA fitness scores (supplemental Figure S2b). The high-GC content of the *M. capsulatus* genome increases the possibility of sgRNA secondary structure formation, which may limit sgRNA targeting efficiency or render some nonfunctional ^56,57^.

We next mapped the sgRNA target sequences from CRISPRi induced and noninduced cultures to the *M. capsulatus* Bath genome to determine overall sgRNA/gene fitness. Given the size of the sgRNA library and multiple sgRNAs per ORF, only those genes with overall positive or negative sgRNA fitness scores were expected to be identified in the analysis. We identified 233 genes whose transcription repression resulted in a fitness defect (sgRNAs depleted in the induced samples, log2 ≤ -1, *p* ≤ 0.001) and repression of 13 genes associated with a fitness advantage (sgRNAs enriched in the induced samples, log2 ≥ 1, *p* ≤ 0.001) (supplemental Figure S3a and supplemental Table S16). Enrichment analysis of the 233 genes with significant sgRNA log2 fitness scores ≤ -1 linked the genes to the essential processes of translation, transcription, ribosome biosynthesis, and amino acid and nucleotide biosynthesis (supplemental Figure S3b and supplemental Table S17), consistent with the analysis using the sgRNA reference library.

The *edd* gene encoding the ED pathway 6-phosphogluconate dehydratase had the most negative differential fitness score (log2 = -3.9, *p* = 1.63e^-71^, Table 3). Several putative essential genes are involved in antioxidant defenses, Fe-S cluster assembly/repair, and disulfide reduction, including superoxide dismutase (MCA2458, log2 = -3.2), a putative Rnf electron transport complex (MCA2893-2899, log2 = -3.1), ferredoxin (MCA2470, log2 = -3.0), and thioredoxin disulfide reductase (MCA1694, log2 = -2.6). These genes may be involved in balancing redox homeostasis during CH_4_ oxidation^58,59^ and represent gene targets for downstream gene-function analysis. sgRNAs targeting genes in the cobalamin biosynthetic operon (MCA2290-MCA2302) had consistent positive fitness scores and MCA2302, encoding a putative CbtB subunit of a cobalt transporter, was identified as significantly enriched (log2 = 1.26, *p* = 3.9e^-11^) when mapping all sgRNAs to the *M. capsulatus* genome (Table 3 and supplemental Table S16). As expected, based on the variable fitness scores for sgRNAs targeting the pMMO and RubisCO operons (supplemental Figure S2), only the *cbbS* (encoding the RubisCO small subunit) and *cbbQ* (encoding a putative RubisCO activase) genes with consistent sgRNA scores were associated with negative fitness scores when mapping to the genome. Interestingly, another putative RubisCO activase gene (MCA2754, log2 = -3.1, *p* = 2.4e^-22^) may be essential for *M. capsulatus* growth on CH_4_. Evaluating the RubisCO activases under differential growth conditions may provide insight into their roles in methanotrophic growth of *M. capsulatus*.

**Table 3.**
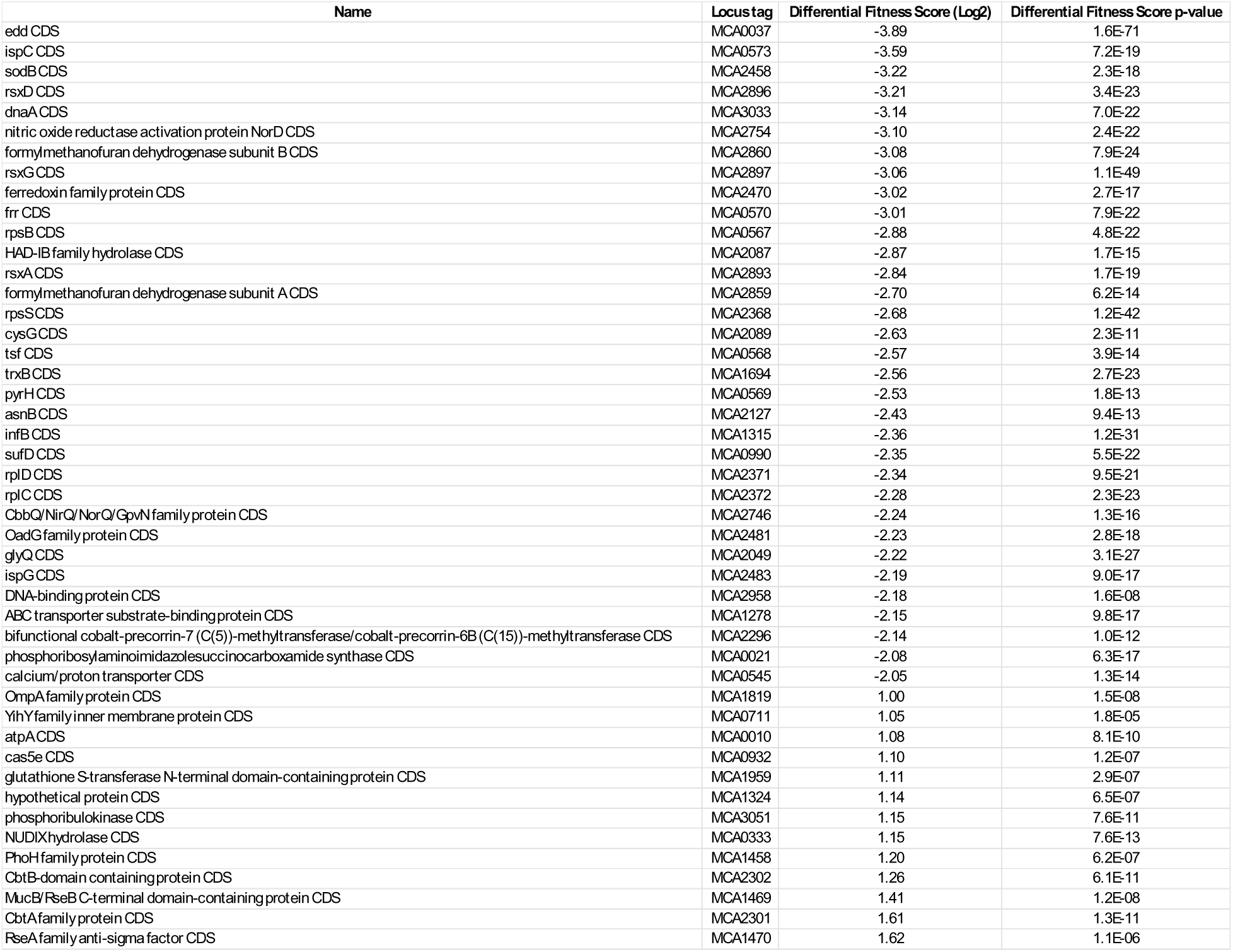
Genes with greatest differential sgRNA fitness scores.

To validate our screening results, we individually targeted five genes with negative fitness scores involved in *M. capsulatus* central metabolism (*edd*, MCA0037, encoding phosphogluconate dehydratase of the Entner-Doudoroff pathway); DNA replication (*dnaA*, MCA3033, encoding the DNA replication initiator), redox metabolism (*fdx*, MCA2470, encoding a ferredoxin; ribosome biogenesis (r*psB*, MCA0567, encoding the S2 protein component of the ribosome 30S subunit); and amino acid biosynthesis (*asnB*, MCA2127, encoding the asparagine synthase); and one gene with a positive fitness score (*cbtB*, MCA2302, encoding a putative cobalt transporter)(Table 3). Consistent with the screening results, high-resolution growth analysis of *M. capsulatus* strains expressing individual sgRNAs displayed significantly inhibited (*edd, dnaA, fdx, rpsB, asnB*) or improved (*cbtB*) growth compared to uninduced controls or a non-targeting sgRNA (Figure 5a-f, Figure 6a). Like that observed when targeting the *cbbL* gene (Figure 3), growth-inhibited strains began to expand after several days of selective pressure. These data validate that the CRISPRi system is effective for functional genomic screening and identification of genes required for CH_4_-dependent growth.

**Figure 5.**
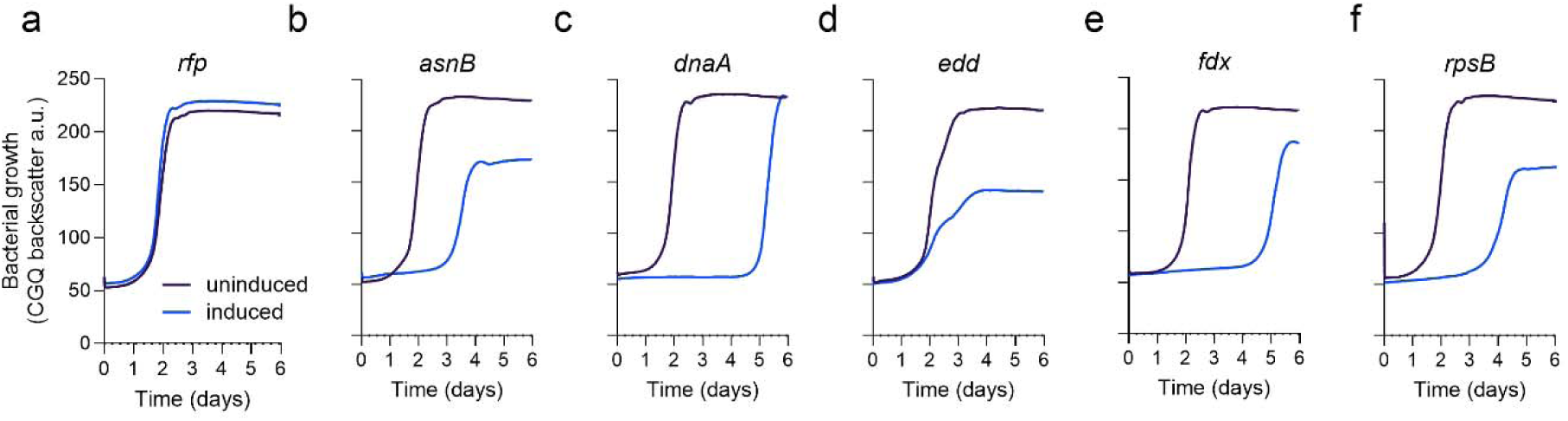
CRISPRi targeted transcription repression of putative essential genes inhibits CH_4_-dependent growth. a-f) High resolution growth curves of strains with sgRNAs targeting genes identified in the genome-wide screen as essential for CH_4_-dependent growth with (blue line) or without (purple line) CRISPRi induction in NMS medium in serum vials with 20% CH_4_ in air atmosphere. A strain expressing an sgRNA targeting mRFP was included as a control (a). Genes represent those encoding enzymes involved in amino acid biosynthesis (b, *asnB*, MCA2127); DNA replication (c, *dnaA*, MCA3033); CH_4_ assimilation (d, *edd*, MCA0037); redox homeostasis (e, *fdx*, MCA2470); and ribosome biosynthesis (f, *rpsB*, MCA0567). Data represent the mean of two independent high-resolution cultivation experiments (n=6).

**Figure 6.**
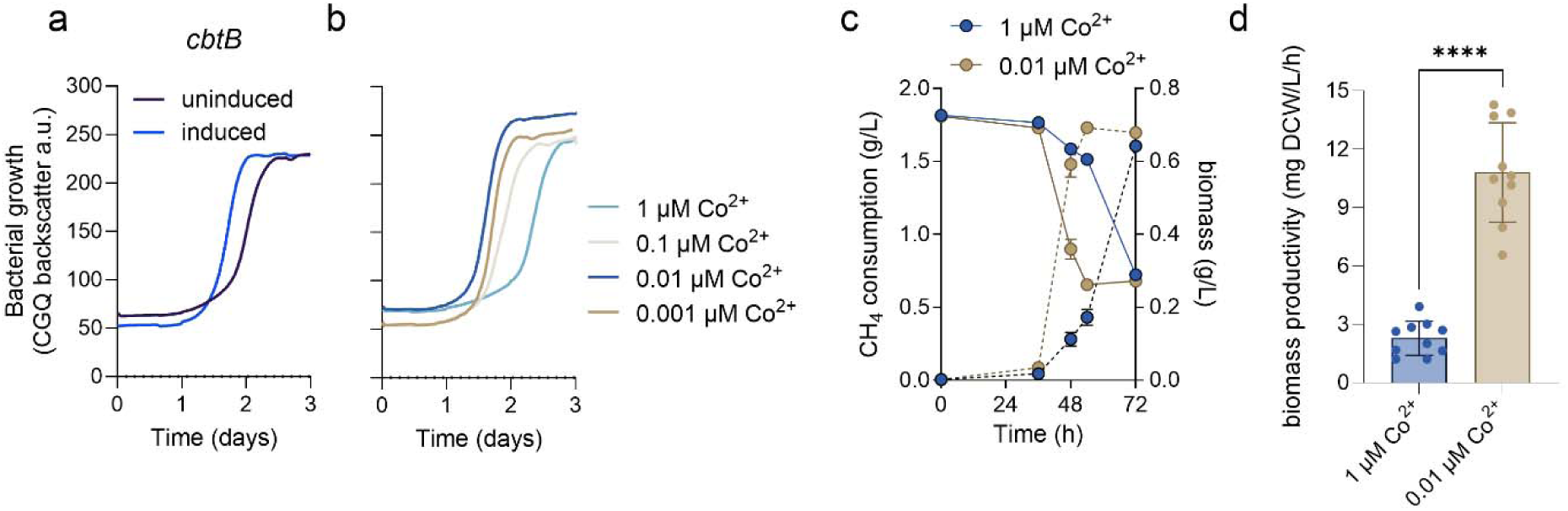
CRISPRi screen-guided growth medium optimization improves methanotroph biomass productivity from CH_4_. High resolution growth curves of a) a strain with an sgRNA targeting the *cbtB* gene with (blue line) or without (purple line) CRISPRi induction or b) wild-type *M. capsulatus* in NMS medium with decreasing CoCl_2_ in the trace elements. c) CH_4_ consumption (solid lines) measured by gas chromatography and biomass accumulation (dashed lines) between wild-type cells cultured in NMS medium with high (1 µM CoCl_2_, blue symbols) or low (0.01 µ CoCl_2_, brown symbols) in serum vials with 20% CH_4_ in air atmosphere. d) *M. capsulatus* biomass productivity in high or low CoCl_2_ NMS medium at 48h. Data in a and b represent the mean of two independent high-resolution cultivation experiments (n=6). Data in c and d represent the mean ± SEM from two independent experiments (n=6-10). **** *p* ≤ 0.0001.

Cobalt is an essential trace element; however, excess can limit bacterial growth by competing with iron and through the generation of reactive oxygen species ^60,61^. Based on the growth improvement observed when transcription of the CbtB cobalt transporter was inhibited (Figure 7a), we hypothesized that the trace amount of cobalt present in our NMS growth medium (1 µM CoCl_2_) was cytotoxic. Consistent with this, *M. capsulatus* cultivated in NMS with decreasing Co^2+^ concentrations displayed a dose-dependent growth improvement, primarily linked to reduced lag phase (Figure 6b). *M. capsulatus* exhibited a 1.4-fold improvement in biomass yield within 48 h in NMS medium with 0.01 µM CoCl_2_ compared to standard NMS (Figure 6c, 0.65 vs 0.48 g DCW/g CH_4_, *p* = 0.006), indicating higher CH_4_ flux to biomass under these conditions. The improved methanotroph growth kinetics in the optimized growth medium equates to a 4.7- fold improvement in methanotroph biomass productivity (Figure 6d). Methanotroph biomass (i.e. single cell protein) is currently generated industrially as a valuable CH_4_-derived product used in the production of animal feed and agricultural fertilizer. Technoeconomic analyses have identified methanotroph productivity as a primary cost driver of methane bioconversion processes^62,63^; thus the optimized medium formulation identified here would be expected to significantly decrease single cell protein production costs.

## Conclusions

Collectively, our results demonstrate that the CRISPRi system developed here can be used for targeted transcription repression in phylogenetically diverse methanotrophic bacteria. Selective screening of a genome-wide sgRNA library during CH_4_-dependent cultivation identified predicted and novel essential genes required for methanotrophic growth. The screen provides genetic evidence that the Entner-Doudoroff pathway is essential for CH_4_ assimilation and the tetrahydromethanopterin pathway is the primary route for CH_4_ dissimilation in *M. capsulatus*. Additionally, the decreased fitness of sgRNAs targeting the ribulose bisphosphate carboxylase/oxygenase (RubisCO) and RubisCO activase genes reiterates the essentiality of CO_2_ assimilation for *M. capsulatus* growth on CH_4_. Notably, our investigations led to the identification of an optimized growth medium for improved methanotroph biomass productivity. The CRISPRi system and associated datasets herein will serve as valuable resources for hypothesis-driven inquiry into genes required for methanotrophy or alternative gene-function analyses. Additionally, these CRISPRi tools can be coupled with other synthetic biology approaches to optimize methanotrophic strains for carbon- and energy-efficient CH_4_ capture and conversion.

## Acknowledgements

This work was supported by a BioDiscovery Institute Seed Grant to CH and RA, the Department of Energy via contract with the MSI STEM Research & Development Consortium (MSRDC) to CH, and the National Science Foundation MCB award # 2225776 to CH. We would like to thank Tallon Coxe in the Azad laboratory for computational support with initial sgRNA library design.

## Author Contributions

CH conceived the project. JH, SL, and YCY performed experiments. CH acquired funding, analyzed and graphed data, and provided overall guidance. CH wrote the manuscript. All others edited and approved the manuscript.

## Data Availability

Plasmids will be available at Addgene upon publication. ONT sequencing data will be deposited at the NCBI Sequence Read Archive (SRA) prior to publication. Code for sgRNA target sequence extraction can be downloaded from the Henard laboratory GitHub database (https://github.com/calvinhenard/CRISPRi-library).

## Supplemental Material

**Figure S1.**
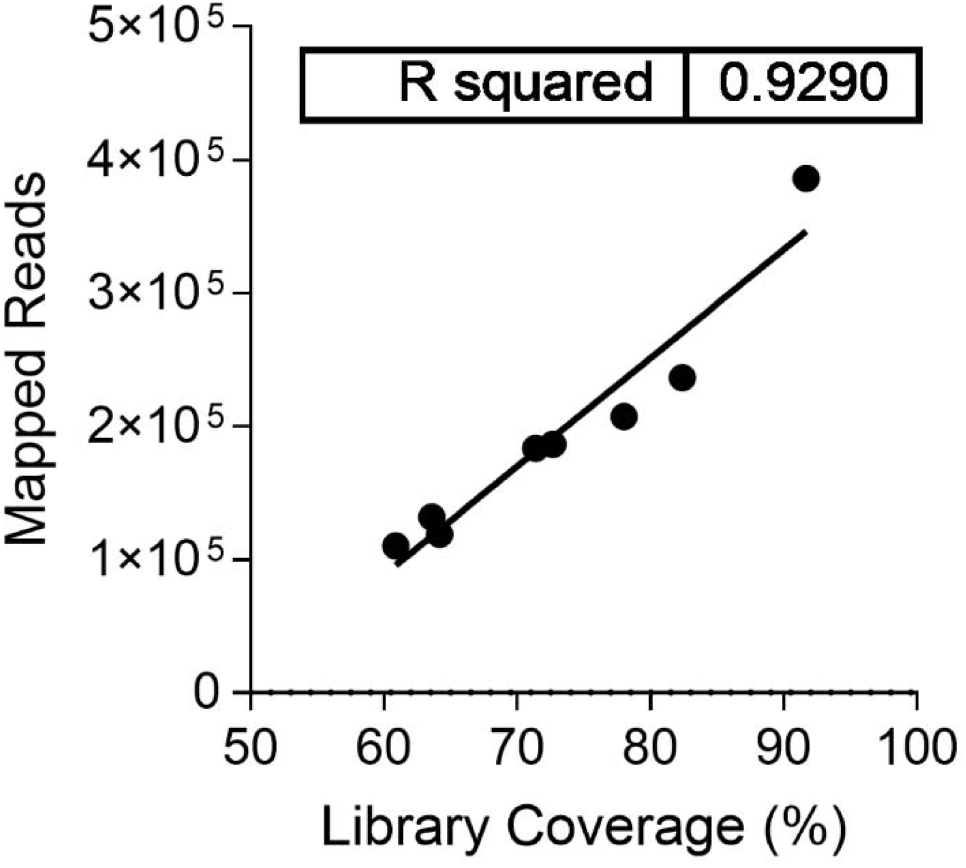
The relationship between sequencing depth and sgRNA library coverage. Oxford Nanopore Technology sequencing reads were processed to extract sgRNA target sequences that were mapped to the synthesized genome-wide sgRNA library. Mapped sequence reads varied between samples. Linear regression analysis suggests 413,960 mapped reads are needed to achieve 100% sgRNA library inclusion in downstream analyses, which is ∼10x sequencing depth.

**Figure S2.**
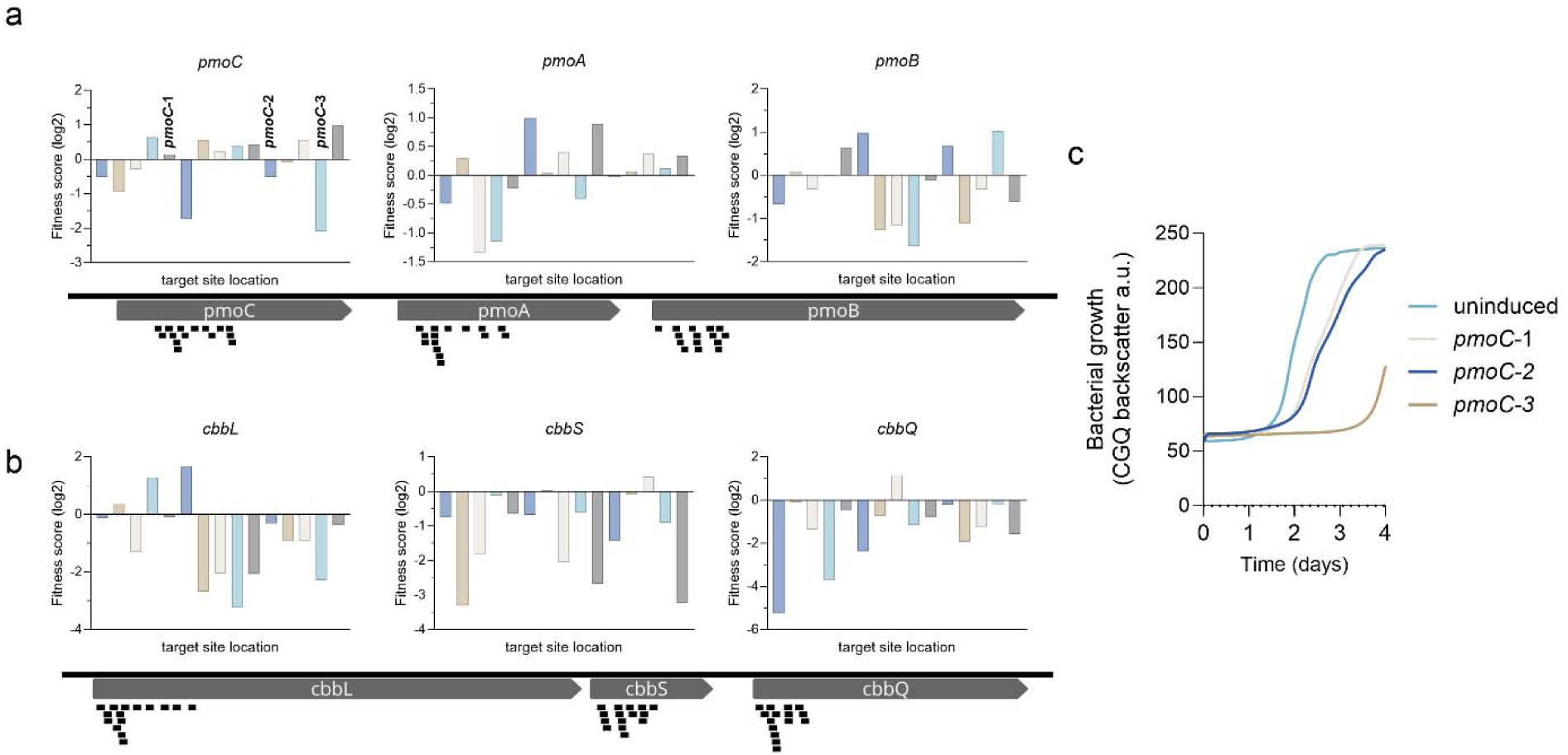
Some sgRNAs targeting the same gene have variable differential fitness scores. Mapped sgRNA target sequences (black rectangles) and corresponding differential fitness score within the *M. capsulatus* a) *pmoCAB* and b) *cbbLSQ* operon genes encoding the particulate methane monooxygenase and ribulose-1,5-bisphosphate carboxylase/oxygenase (RubisCO) and associated RubisCO activase, respectively. c) High resolution CH_4_-dependent growth curves of strains with sgRNAs with differential fitness scores (*pmoC-1*, *pmoC-2*, *pmoC-3*) targeting the *pmoC* gene with (blue line) or without (purple line) CRISPRi induction. The growth curves represent the mean of 4-6 independent observations. The mean of the three uninduced sgRNA controls is shown.

**Figure S3.**
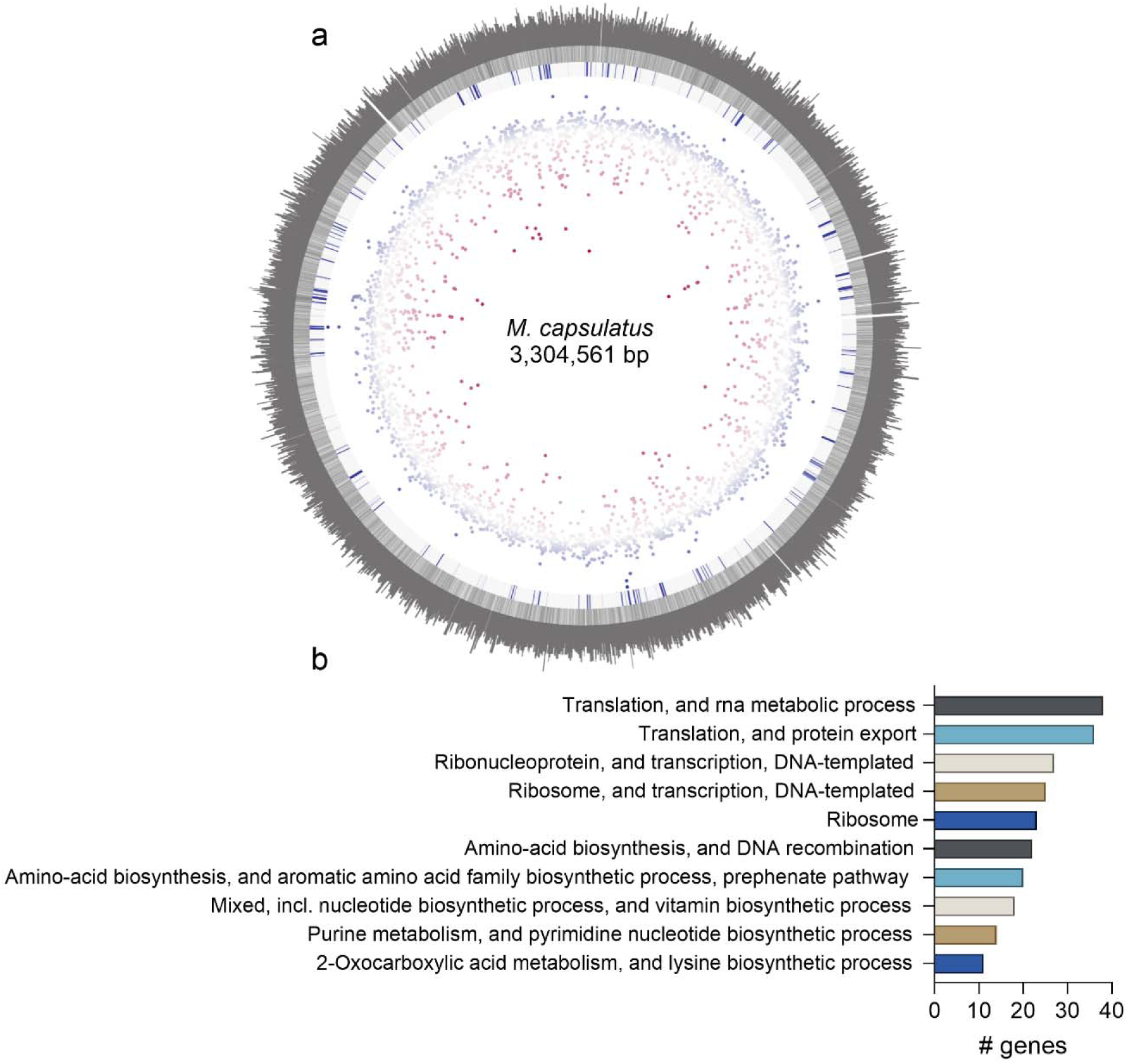
A CRISPRi genome-wide screen identifies genes essential for CH_4_-dependent growth of *M. capsulatus*: mapping to the *M. capsulatus* genome. a) Circos plot of the methanotroph sgRNA library raw reads (outer ring) mapped to the NC_002977 *M. capsulatus* reference genome (2^nd^ ring, genes in forward direction are in dark grey and reverse direction in light grey). Differential sgRNA fitness scores determined via DeSeq2 between CRISPRi induced and uninduced samples during selection/cultivation with CH_4_ as the sole carbon and energy source (4^th^ ring, blue dots log2 >1; red dots log2 <1). sgRNAs depleted in the screen (negative fitness scores) correspond to genes with transcription repression causing a growth defect where those enriched (positive fitness scores) correspond to genes with transcription repression causing a growth advantage. sgRNAs with statistically significant (*p* ≤ 0.01) differential fitness scores are indicated (blue squares, 3^rd^ ring). b) ShinyGo enrichment analysis of the 233 genes with sgRNA log2 fitness scores ≤ -1 and *p* ≤ 0.001.

Supplemental Tables S1-S17 can be found here: https://docs.google.com/spreadsheets/d/1I1Mwm8o_4b5vcYRpeE11nL-bj-knuRPL/edit?usp=drive_link&ouid=103067052205615496775&rtpof=true&sd=true

